# Sodium/Potassium ATPase Alpha 1 Subunit Fine-tunes Platelet GPCR Signaling Function and is Essential for Thrombosis

**DOI:** 10.1101/2024.05.13.593923

**Authors:** Oliver Q. Li, Hong Yue, Autumn R. DeHart, Renat Roytenberg, Rodrigo Aguilar, Olalekan Olanipekun, Fang Bai, Jiang Liu, Olga Fedorova, David Kennedy, Ellen Thompson, Sandrine V. Pierre, Wei Li

**Author notes:** Corresponding author: Wei Li, MD, PhD, FAHA Department of Biomedical Sciences, BBSC 241G Joan C. Edwards School of Medicine Marshall University One John Marshall Drive, Huntington, West Virginia 25755-9310, USA. Or Sandrine V. Pierre, PhD Marshall Institute for Interdisciplinary Research Department of Biomedical Sciences, Joan C. Edwards School of Medicine Marshall University One John Marshall Drive, Huntington, West Virginia 25755-9310, USA.

## Abstract

**Background:** Thrombosis is a major cause of myocardial infarction and ischemic stroke. The sodium/potassium ATPase (NKA), comprising α and β subunits, is crucial in maintaining intracellular sodium and potassium gradients. However, the role of NKA in platelet function and thrombosis remains unclear.

**Methods:** We utilized wild-type (WT, α*1^+/+^*) and NKA α1 heterozygous (α*1^+/-^*) mice, aged 8 to 16 weeks, of both sexes. An intravital microscopy-based, FeCl_3_-induced carotid artery injury thrombosis model was employed for in vivo thrombosis assessment. Platelet transfusion assays were used to evaluate platelet NKA α1 function on thrombosis. Human platelets isolated from healthy donors and heart failure patients treated with/without digoxin were used for platelet function and signaling assay. Complementary molecular approaches were used for mechanistic studies.

**Results:** NKA α1 haplodeficiency significantly reduced its expression on platelets without affecting sodium homeostasis. It significantly inhibited 7.5% FeCl_3_-induced thrombosis in male but not female mice without disturbing hemostasis. Transfusion of α*1^+/-^*, but not α*1^+/+^*, platelets to thrombocytopenic WT mice substantially prolonged thrombosis. Treating WT mice with low-dose ouabain or marinobufagenin, both binding NKA α1 and inhibiting its ion-transporting function, markedly inhibited thrombosis in vivo. NKA α1 formed complexes with leucine-glycine-leucine (LGL)-containing platelet receptors, including P2Y12, PAR4, and thromboxane A2 receptor. This binding was significantly attenuated by LGL>SFT mutation or LGL peptide. Haplodeficiency of NKA α1 in mice or ouabain treatment of human platelets notably inhibited ADP-induced platelet aggregation. While not affecting 10% FeCl_3_-induced thrombosis, NKA α1 haplodeficiency significantly prolonged thrombosis time in mice treated with an ineffective dose of clopidogrel.

**Conclusion:** NKA α1 plays an essential role in enhancing platelet activation through binding to LGL-containing platelet GPCRs. NKA α1 haplodeficiency or inhibition with low-dose ouabain and marinobufagenin significantly inhibited thrombosis and sensitized clopidogrel’s anti-thrombotic effect. Targeting NKA α1 emerges as a promising antiplatelet and antithrombotic therapeutic strategy.

## Introduction

Thrombotic events represent a substantial global health concern, contributing significantly to morbidity and mortality worldwide^1,2^. Platelet activation and hyper-aggregation at vascular injury sites constitute the primary pathogenic processes underlying thrombosis, leading to vessel occlusion, myocardial infarction, and stroke. Platelet surface glycoproteins (GP), such as GPIb-IX-V complex, GPIIb/IIIa (integrin αIIbβ3), and GPVI, play crucial roles in mediating both platelet adhesion and activation in response to exposed agonists at the site of vascular injury^3–7^. Activated platelets release soluble agonists, including adenosine diphosphate (ADP), thrombin, and thromboxane A2 (TXA2)^8,9^, which locally activate additional platelets via G protein-coupled receptors (GPCRs), such as the ADP receptor P2Y12 and thrombin receptor protease-activated receptor-1 (PAR1)^6,10^. Current clinical strategies for preventing platelet-mediated thrombotic events involve antiplatelet drugs, such as aspirin [Cyclooxygenase inhibitor, reduces TXA2 expression and thus reduces TXA2 receptor (TP) mediated platelet activation], clopidogrel (P2Y12 inhibitor), and vorapaxar (PAR1 inhibitor)^11–14^. However, these drugs often come with major side effects, such as systemic or gastrointestinal hemorrhage^2,11–15^. Some patients also remain unresponsive to these regimens, experiencing a high incidence of vascular thrombosis^2^. Certain medical conditions, such as heart failure (HF), elevate the risk of thrombosis, yet the underlying mechanisms remain unclear. Therefore, there exists a critical need to explore novel molecular mechanisms of platelet activation. This exploration will inspire the development of innovative antiplatelet therapies, aiming to reduce and prevent thrombosis with minimal risk of systemic side effects.

Platelets contain numerous proteins, and many of their functions, particularly in the context of thrombosis, remain unknown. The sodium/potassium ATPase (NKA), also known as sodium pump, is a transmembrane protein composed of α and β subunits^16^. Currently, four α and three β isoforms have been identified, forming various NKA isoenzymes in a tissue- and cell-specific manner^17–19^. Based on three platelet-proteomic studies^20–22^ and previous pioneering work, the platelet NKA is presumed to be α1β3. NKA plays a pivotal role in maintaining intracellular Na^+^ and K^+^ concentrations, transmembrane electrical gradient, and cell volume. The NKA α1 subunit, encoded by *Atp1a1*, also serves as a signal transducer, influencing cell growth and differentiation^17,23–25^. The role of NKA in platelet function and thrombosis remains nebulous, with published data presenting conflicting findings^26^. A systematic study of NKA’s role in platelet biology and thrombosis is lacking.

NKA is highly sensitive to inhibition by cardiotonic steroids (CTS), most notably the digitalis class, which has been used widely in the treatment of HF and arrhythmias^23,27^. Scattered studies have suggested that inhibiting NKA pump function with ouabain enhances platelet aggregation in vitro^28–30^, mediated by the increase of intracellular Ca^2+^-evoked exposure of phosphatidylserine^47^. Several studies have also suggested that digoxin usage increases the risk of thrombotic events^31,32^. Patients with HF exhibit adverse platelet characteristics, including reduced survival time, increased mean platelet volume, and heightened activation and reactivity, impacting their morbidity and mortality^33,34^. However, the underlying mechanism for both HF- and CTS-associated high-risk of thrombosis is unclear. Ligands of platelet GPCRs, such as thrombin, ADP, and TXA2, have been reported to inhibit the NKA pump function^35,36^, suggesting a potential crosstalk between NKA and certain platelet GPCRs. However, the underlying molecular mechanisms and the pathophysiological significance of these phenomena are also not known. In this study, we aimed to explore a novel function of NKA α1, elucidating how CTS regulate platelet activation and thrombosis.

## Methods Animals

The α*1^+/-^* mouse strain was generated by Dr. Jerry Lingrel at the University of Cincinnati College of Medicine and has been backcrossed with the C57BL6/J wild-type (WT) mice more than 10 times^37^. The mouse strain was kept in heterozygous inbreeding and WT littermates (α*1^+/+^*) were used as controls. All procedures and manipulations of animals have been approved by the Institutional Animal Care and Use Committee of Marshall University (#: 1033528, PI: WL).

### Materials

Platelet agonists including ADP (P/N 384), collagen (P/N 385), and thrombin (P/N 386) were purchased from Chrono-log (Havertown, PA). Collagen-related peptide (CRP) was a gift from Dr. Peter Newman (Blood Research Institute, WI)^38^. Antibodies to phosphorylated AKT (4060S), pan-AKT (2920S), Phospho-Src Family (Tyr416) (E6G4R, Rabbit mAb), Src (36D10, Rabbit mAb), and HRP-conjugated Pan-actin (12748S), HRP Conjugated anti-Rabbit IgG Light-Chain Specific secondary antibody (D4W3E, #93702), and HRP Conjugated anti-mouse IgG Light-Chain Specific secondary antibody (D2V2A, #58802) were purchased from Cell Signaling Technology (Danvers, MA). Antibodies to P2Y12 (AGP-098) and A2BR (AAR-003) were purchased from Alomone Lab. Antibodies to NKAα1 subunit (ab2872, ab7671, and ab76020) and P2Y12 (ab184411, ab183066) were purchased from abcam (Waltham, MA). Antibody to NKAα1 subunit (clone C464.6, Cat# 05-369) and mouse anti-Flag antibody (F1804) was purchased from Millipore-Sigma. Antibodies to TP (27159-1-AP), P2Y1 (67654-1-AP), and PAR4 (25306-1-AP) were purchased from Proteintech Group, Inc (Rosemont, IL). Ouabain and marinobufagenin (MBG) were purchased from Cayman Chemical (Ann Arbor, MI). The Polyvinylidene difluoride (PVDF) membrane was purchased from Bio-Rad. The Novex 3-12% NativePAGE Bis-Tris gel, NativePAGE sample prep kit, NativeMark unstained protein standard, NativePAGE running buffer, NativePAGE cathode buffer additive, and dithiothreitol (DTT) were purchased from ThermoFisher Scientific Invitrogen (Calsbad, CA). All other chemical reagents were purchased from Millipore Sigma (Burlington, MA) except where specifically indicated.

### Murine FeCl_3_-injury-induced carotid artery thrombosis model and tail bleeding assay

The ferric chloride (FeCl_3_)-injury-induced carotid artery thrombosis model has been described previously^39,40^. In brief, mice in both sexes, 8 to 16 weeks old, were anesthetized by a mixture of ketamine/xylazine (100/10 mg/kg) via intraperitoneal injection. The right jugular vein was exposed and injected with 100 μl of rhodamine 6G solution (Sigma 252433-1G, 0.5 mg/ml in saline, 0.2 μm filtered) to label platelets. The carotid artery injury was induced by topically applying a piece of filter paper (1 x 2 mm) saturated with 7.5% or 10% FeCl_3_ solution for 1 minute. Thrombi formation was observed in real-time using intravital microscopy with a Leica DM6 FS fluorescent microscope (Deerfield, IL, USA) attached to a Gibraltar Platform. Video imaging was conducted using a QImaging Retiga R1: 1.4 Megapixel Color CCD camera system with mono color mode (Teledyne Photometrics, Tucson, AZ, USA) and StreamPix version 7.1 software (Norpix, Montreal, Canada). The endpoints were set as 1) blood flow has ceased for > 30 seconds, or 2) occlusion is not seen after 30 minutes of FeCl_3_ injury. In the second case, 30 minutes were assigned to the mouse for statistical analysis.

### Tail bleeding assay

The tail bleeding assay was conducted on the mice following the thrombosis study by truncating the mouse tail 1 cm from the tip. The truncated tail was immediately immersed in 37°C warm saline, and the time from tail truncation to the cessation of blood flow was recorded^38,41,42^.

### Murine platelet isolation

Mice were anesthetized with ketamine/xylazine (100/10 mg/kg), and 0.9 – 1 mLwhole blood was collected through inferior vena cava puncture using 0.109 M sodium citrate as an anticoagulant^42,43^. Modified Tyrode’s buffer (concentration of components in mM: 137 NaCl, 2.7 KCl, 12 NaHCO_3_, 0.4 NaH_2_PO_4_, 5 HEPES, 0.1% glucose, and 0.35% BSA, pH 7.2) was added to the collected whole blood at 0.7 volumes, mixed, and then platelet-rich plasma (PRP) was isolated by centrifugation at 100 g for 10 min. The sedation containing red blood cells and leukocytes was further centrifuged at 13,000 rpm for 1 min to isolate platelet-poor plasma (PPP).

For the preparation of washed platelets, a final concentration of 0.5 µM PGE1 was added to the PRP, followed by centrifugation at 650 g for 6 min to pellet platelets. Platelets were re-suspended in PBS containing 0.109 M sodium citrate (at a ratio of 9:1) and 0.5 µM PGE1, and then re-pelleted (650 g, 6 min). Finally, platelets were resuspended in PBS, counted with a hemocytometer, and used immediately.

### Platelet aggregation assay

The platelet concentration in PRP was counted with a hemocytometer and adjusted to 2.5E+08/ml with PPP, and 0.4 ml of this platelet suspension was used for the platelet aggregation assay using Chrono-log 700 with an agitation speed of 1,200 rpm. CaCl_2_/MgCl_2_ was repleted at a final concentration of 1 mM immediately before adding a platelet agonist^38,42,43^.

### Cellix flow chamber-based platelet adhesion and aggregation assay

Cellix flow chambers were coated overnight with 10 µL 100 μg/mL collagen in PBS, as previously mentioned^42,44,45^. Whole blood was drawn from the α*1^+/-^*and α*1^+/+^* male mice via the inferior vena cava. The whole blood was labeled with Rhodamine 6G (at a final concentration of 50 µg/ml), repleted with CaCl_2_/MgCl_2_ to a final concentration of 1 mM, and then perfused through the flow chamber at a shear rate of 65 dyne/cm^2^ for 3 minutes. The chamber was rinsed with PBS at the same flow rate for 3 minutes, and images were captured at the 6 marker positions as specified by the manufacturer. The fluorescent intensity of each image was analyzed with Image J software. Data were presented as the mean value of each image minus the mean value of the background (without cells) in the same image.

### Platelet sodium-potassium ATPase pumping function assay

Platelets in PRP isolated from the α*1^+/-^* and α*1^+/+^*male mice were loaded with the ^22^Na^+^ at a concentration of 1 µCi/ml medium for 1 hour at 37°C. After 4 rapid washes with PBS, 500 uL of 1N NaOH was applied to the platelet pellet and incubated at room temperature for 30 minutes to extract platelet-associated ^22^Na^+^. Radioactivity in the extraction solution was assessed using a scintillator, and the data were presented as counts per minute (CPM). The platelets were then lysed with 1N 0.1N NaOH/0.2% SDS at room temperature for 30 min and protein concentration was measured. The NKA activity was calculated as ^22^Na^+^ CPM count corrected by total protein mass in micrograms^46^.

### Evaluation of P2Y12-mediated platelet signaling activation

Platelets in PRP (2.5E+08/ml) were stimulated with 2.5 μM ADP for 0, 1, 3, and 5 min^38^. Platelet activation was stopped by adding 1 mM EDTA and 0.5 µM PGE1 to the reaction mixture. The platelets were then pelleted by centrifugation at 13,000 RPM for 15 seconds, lysed in radioimmunoprecipitation assay (RIPA) buffer containing Halt™ Protease and Phosphatase Inhibitor Single-Use Cocktail, EDTA-Free (100 X) (ThermoFisher, Cat# 78443). Protein concentration was determined using a Bio-Rad Protein assay. AKT activation was assessed by immunoblotting assays of AKT phosphorylation.

### Western blot assay of platelet **α1** expression

Washed platelets isolated from α*1^+/-^* and α*1^+/+^* male mice were lysed in RIPA buffer containing the proteinase and phosphatase cocktail. Thirty micrograms of total protein were used for the western blot assay of α1 expression. Actin was re-blotted in the same membrane as a loading control. All samples involving western blot assay of α1 were mixed with 2 X Laemmli sample buffer containing 2-mercaptoethanol (except non-reduced condition) in 1:1 volume ratio and then incubated at 37 °C for 30 min before loading to the gel.

### Platelet transfusion studies

α*1^+/+^*mice were exposed to 11 Gy of irradiation to induce thrombocytopenia, with platelet counts falling below 5% of normal levels after 5 days^47^. Donor platelets were isolated from male α*1^+/-^* and α*1^+/+^* mice and stained with Rhodamine 6G (final concentration 50µg/ml) for 5 min at room temperature. Platelets isolated from one donor mouse (about 1 mL whole blood) in 150 µL saline were injected into one thrombocytopenic recipient mouse through the jugular vein, and then the thrombosis study was initiated 10 minutes later^38^.

### Sodium/Potassium ATPase inhibition on human platelet aggregation

Whole blood was collected from healthy human volunteers under the approval of the Marshall University Institutional Review Board (IRB#1185274 and IRB#1347920, PI: Wei Li). PRP was isolated by centrifugation in a swing bucket centrifuge at 170 g for 15 minutes. The top 2/3 of the PRP was transferred into a new tube, and the remaining portion was further centrifuged at 2,000 g for 20 minutes to isolate PPP. The platelet concentration was adjusted to 2.5E+08/ml with PPP, and 0.4 ml of this suspension was used for platelet aggregation assay with an agitation speed of 1,200 rpm. To test the role of NKA inhibition on platelet aggregation, ouabain was added into the reaction solution to a final concentration of 0, 25, 50, and 100 nM. The solution was then incubated for 5 min in the cuvettes, after which aggregation was initiated by repleting CaCl_2_/MgCl_2_ to a final concentration at 1 mM and ADP to a final concentration of 2.5 µM.

### Immunoprecipitation (IP) and Immunoblotting (IB) assays

One thousand micrograms of human and mouse platelet lysates prepared as mentioned above, as well as COS-7 cell lysates (prepared as mentioned below), were used for standard IP-IB assays using specific antibodies as indicated in the Results section. Briefly, samples in 1 mL RIPA or IP lysis buffer were pre-cleaned by incubating with 450 µg Dynabeads™ Protein G (Cat# 10004D, ThermoFisher Scientific) with gentle rotation for two hours at 4 °C. Subsequently, 2 µg of the specific antibody was added into the reaction solution and incubated overnight with gentle rotation. The immune complex was pulled down by adding 1.5 mg Dynabeads™ Protein G and incubated at 4 °C with gentle rotation for two hours. The beads were washed three 15 minutes with RIPA or IP lysis buffer and then eluted with 50 µL 2 x Laemmli buffer. Twenty-five micrograms of input samples were run in the same gel as controls for identifying positive signals.

### Identifying the NKAα1 associated platelet GPCR

To investigate how α1 affects platelet GPCR signaling function and its interaction with certain platelet GPCRs, we transfected COS-7 cells (ATCC, CRL-1651) with plasmids encoding P2Y12 (item#66471, Addgene), F2RL3 (PAR4) (item #66279), TP (item #66517), or A2BR (item#37202) using Lipofectamine™ 3000 Transfection Reagent (L3000015, ThermoFisher Scientific). After 36 hours, cells were lysed and the lysates were used for co-IP assay to assess the interaction between NKAα1 (endogenous) and these GPCRs.

The binding of PAR4 and P2Y12 in human platelets is mediated by a peptide containing three amino acids, leucine-glycine-leucine (LGL)^48^. Both human PAR4 and P2Y12 contain the LGL motif. This LGL motif is conserved in mouse P2Y12 (amino acid 121-123). Human α1 also contains an LGL motif (884-886), and the corresponding sequence in the mouse is leucine-glycine-isoleucine (LGI). To determine if this LGL(LGI) motif mediates the α1 and P2Y12 binding, we constructed LGL(LGI)> serine-phenylalanine-threonine (SFT) mutant murine NKAα1 and P2Y12. To this end, α*1* (encoded by *Atp1a1*) gene was amplified from pLV[Exp]-EGFP:T2A:Puro-CMV>*mAtp1a1*[NM<u>_</u>144900.2] (VectorBuilder, Chicago, IL) using Invitrogen Platinum SuperFi II PCR Master Mix (2X) with primers as shown in the **Table 1**. The PCR product was purified and cloned into a pCDNA6B vector using HindIII and EcoRV cloning sites.

**Table 1.**
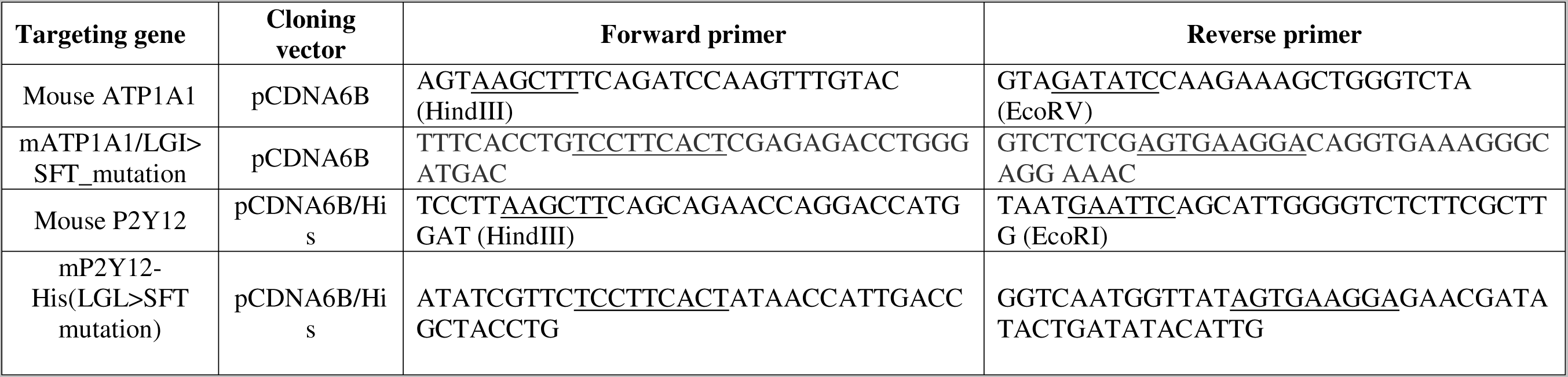
List of primers for amplifying the genes targeted.

Mouse brain cDNA was used as a template to amplify mouse P2Y12 using primers listed in Table 1, and subsequently cloned into pCDNA6B using HindIII and EcoRI cloning sites. This cloning generated a His-tagged mouse P2Y12. The plasmid pCDNA6B/mP2Y12-His and pCDNA6B/*mAtp1a1* were used as a template and the LGL motif in the mouse P2Y12 and the LGI motif in the mouse NKAα1 was mutated to SFT using PCR-directed point mutation. The whole plasmid was sequenced and the mutations were confirmed using the service provided by the “plasmidsaurus”. These plasmids were co-transfected into COS-7 cells using the composition as indicated in the Results. Cells were harvested 36 hours post-transfection and used for IP-IB assays to determine how LGL>SFT mutation in P2Y12 or LGI>SFT mutation in mouse NKAα1 affects the interaction of α1 and P2Y12.

### Blue native polyacrylamide gel electrophoresis assay for identifying the interaction between ATP1A1 and P2Y12

The sample preparation and analysis for Blue Native Polyacrylamide Gel Electrophoresis (BN-PAGE) followed the previous publication^43,49^. Briefly, platelets isolated from the α*1^+/-^*and α*1^+/+^* male mice were lysed with 1X BN-PAGE sample buffer (with 2% DDM, n-dodecyl-β-D-maltoside and Halt Protease and Phosphatase Inhibitor Cocktail) and cleared by centrifugation at 20,000 x g for 30 min at 4°C and stored in -80°C until use. Immediately before sample loading, the NativePAGE 5% G-250 sample additive was added to samples (at 1/4th of detergent DDM concentration) and mixed. Electrophoresis was conducted according to the manufacturer’s instructions. Proteins dissolved in the gel were transferred onto the PVDF membrane and followed standard immunoblotting procedures using antibodies against α1 and P2Y12.

### Effect of cardiotonic steroids on thrombosis

To determine if CTS treatment mimics the α1 deficiency in mice, we treated WT C57BL/6J mice with ouabain (100 µg/Kg) and MBG (100 µg/Kg)^50^ and then subjected these mice to the thrombosis study around 18 hours after drug administration using the 7.5% FeCl_3_-injury-induced thrombosis model.

### To determine if NKAα1 deficiency or inhibition could reduce the effective doses of the current antiplatelet drugs and thus attenuate side effects

To determine if α*1* deficiency mediated anti-thrombotic effect could benefit the dual antiplatelet therapy, we treated α*1^+/-^* and α*1^+/+^*male mice with 0.5 and 1 mg/kg Clopidogrel, which has been demonstrated to have no anti-thrombotic effect on WT mice^42^, by gavage feeding for one week and then subjected to the thrombosis study using 10% FeCl_3_-injury-induced thrombosis model.

### Statistics

Data are expressed as mean ± SEM. Results were analyzed by 2-tailed Student’s *t* test, Mann Whitney test, or 1-way ANOVA with Bonferroni post-hoc test for multiple comparisons using GraphPad Prism (version 10.2.2). In some cases, data were analyzed by Log-rank test using the Kaplan-Meier survival curve. *P* < 0.05 was considered statistically significant.

## Results

### Alpha 1 gene haplodeficiency results in a significant anti-thrombotic effect in male mice without affecting bleeding

The α*1^+/-^* mouse strain was generated by Dr. Jerry Lingrel at the University of Cincinnati College of Medicine and has been backcrossed with the C57BL6/J wild-type (WT) mice more than 10 times^37^. To determine if NKA α1 affects thrombosis, we conducted an in vivo thrombosis study in α*1*^+/-^ mice using the 7.5% FeCl_3_-injury-induced carotid artery thrombosis model. As shown in **Fig. 1A** and **1B** as well as online video 1, thrombus formation in the male α*1*^+/+^ mice was the same as seen in the C57BL6/J WT male mice with an average thrombosis time of around 11 min^51,52^. Alpha 1 haplodeficiency in male mice did not significantly affect initial platelet adhesion and aggregation at the site of the injured vessel wall (Fig. 1A, 1 min after injury); however, the second phase of platelet activation, which is mainly mediated by platelet GPCRs and the activation of the coagulation system^53,54^, was dramatically attenuated, resulting in a significantly prolonged time to form occlusive thrombosis in male α*1*^+/-^ mice (Fig. 1A, 1B and online video 2). In most of the α*1*^+/-^ mice, a large thrombus was formed at the end of the 30-minute observation, but without blood flow cessation. Histological examination of the carotid artery with thrombi collected from the middle of the injured vessel confirmed the presence of holes in the thrombi harvested from the α1^+/-^ mice, but not from the α*1*^+/+^ mice (Supplemental Figure 1). However, α*1* haplodeficiency did not affect thrombosis in female mice (*p* = 0.39, Fig. 1B). Consequently, in the following studies, we only used male mice to clarify the role of NKA α1 in platelet activation and thrombosis.

**Fig. 1.**
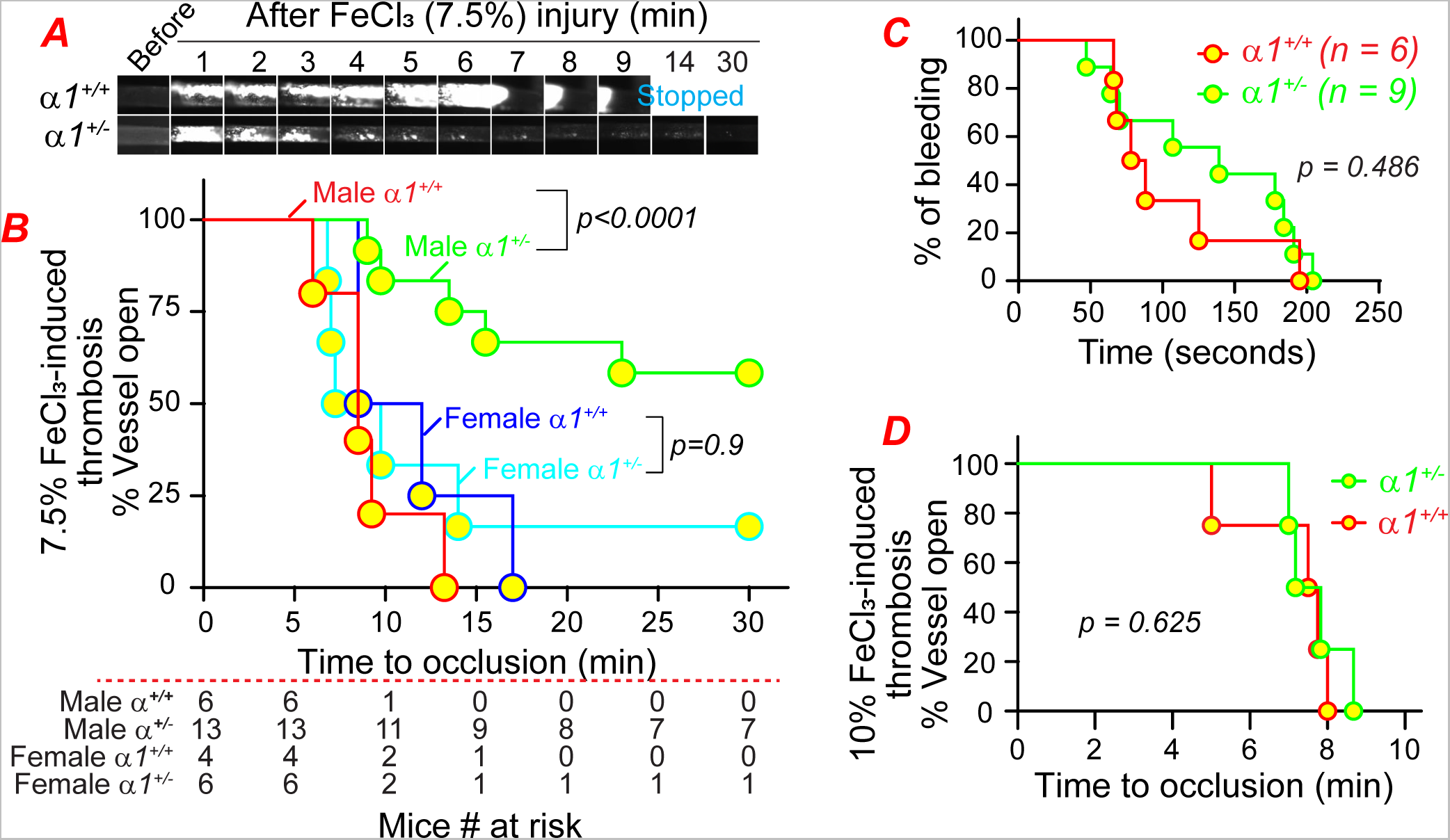
Alpha1 haplodeficiency attenuates arterial thrombosis in male mice without affecting hemostasis. ***A.*** Representative video images: Thrombus formation in the carotid artery was induced by 7.5% FeCl_3_. Platelets were labeled via direct intravenous injection of Rhodamine 6G solution in saline. ***B.*** Accumulated data: Time to occlusive thrombus formation after 7.5% FeCl_3_ injury was assessed. ***C.*** Tail bleeding assay was conducted by truncating the tail 1 cm from the tip. ***D.*** Thrombus formation in the carotid artery was induced by 10% FeCl_3_ treatment in male mice. The logrank (Mantel-Cox) test was used for the statistical analyses in panels B, C, and D.

Despite the significantly prolonged thrombosis time, tail bleeding time was the same between the male α*1^+/+^* and α*1^+/-^* mice (**Fig. 1C**). Furthermore, under a more severe injury induced by 10% FeCl_3_, the time to carotid artery occlusion in male mice became similar between the two strains (**Fig. 1D**). These findings suggest that NKA α1 plays a mechanistic role during platelet activation and thrombosis, while sex hormones may also contribute to this context.

### Platelet NKA α1 expression accounts for the anti-thrombotic phenotype in male α*1^+/-^* mice

To clarify how NKA α1 affects thrombosis, we first examined α1 expression in platelets. As shown in **Fig. 2A**, α*1* gene haplodeficiency dramatically reduced α1 protein expression in platelets harvested from the male mice. However, α1 haplodeficiency did not affect intracellular sodium levels (**Fig. 2B**). Therefore, the observed antithrombotic effect in α*1*^+/-^ male mice is unlikely to be attributable to the increase of Ca^2+^-evoked exposure of phosphatidylserine^47^.

**Fig. 2.**
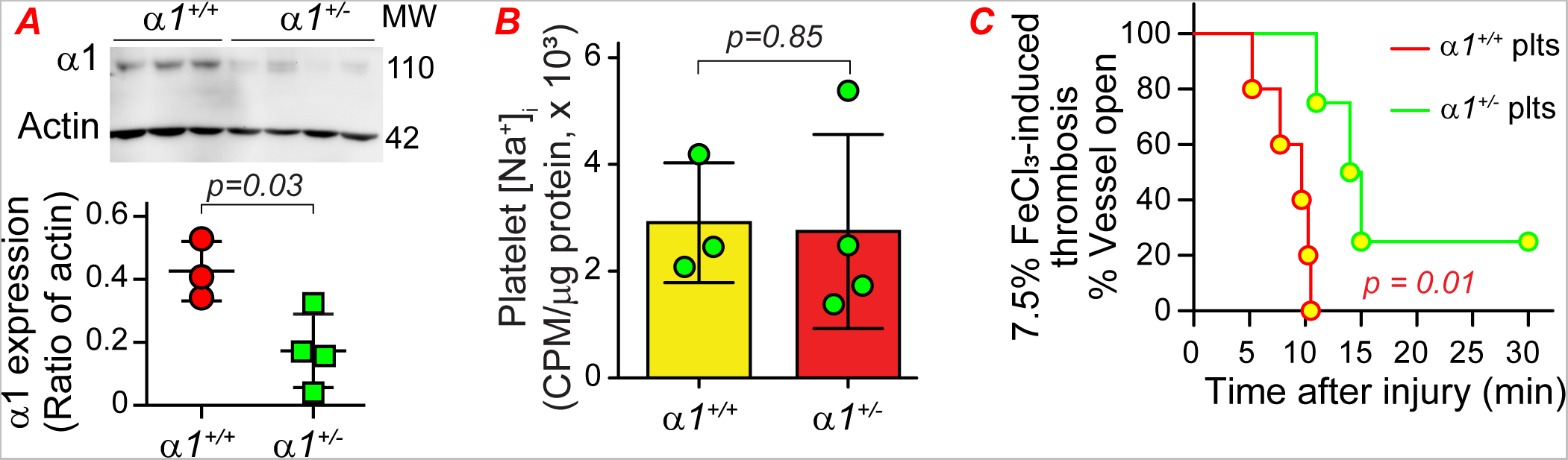
Platelet α1 is essential for normal platelet function and thrombosis. ***A.*** Platelets isolated from male α*1^+/-^* and α*1^+/+^* mice were used for western blot assay of α1 expression. ***B.*** Intracellular sodium concentration was assessed in platelets isolated from male α*1^+/-^* and α*1^+/+^* mice. ***C.*** Male WT mice were lethally irradiated with X-RAD320 at a dose of 11 Gy. Five days later, these mice were infused with platelets isolated from the male α*1^+/-^* and α*1^+/+^* mice via jugular vein injection, and then thrombosis was initiated using 7.5% FeCl_3_ injury 10 min later.

We hypothesized that the anti-thrombotic phenotype found in the male α*1^+/-^* mice was due to the decreased α1 expression in platelets, but not due to the reduction of α1 in the vessel wall cells or other circulating cells. To test this hypothesis, we conducted a platelet transfusion assay as reported in previous studies^38^. As shown in **Fig. 2C**, compared to thrombocytopenic WT recipients that received α*1^+/+^* platelets, thrombocytopenic WT mice that received α*1^+/-^* platelets had a dramatically prolonged time to form occlusive thrombosis. These data suggest that platelet α1 is responsible for the antithrombotic phenomena found in the α*1^+/-^* mouse strain.

### Alpha1 haplodeficiency significantly reduces ADP-induced platelet aggregation

To understand how α1 affects platelet activation and thus thrombosis, we conducted a Cellix flow chamber assay and examined how α*1^+/-^* and α*1^+/+^* platelets adhere to collagen-coated surfaces. As shown in **Fig. 3A**, α1 deficiency did not affect platelet adhesion to the collagen-coated surface, suggesting that NKA α1 does not affect collagen receptor-mediated platelet adhesion and activation. We then conducted a platelet aggregation assay and tested how α*1^+/-^*platelets in platelet-rich plasma (PRP) respond to the conventional platelet agonists. As shown in **Fig. 3B**, in line with the finding of Cellix Flow Chamber assay, collagen (1 µg/ml) induced platelet aggregation was the same between the α*1^+/-^* and α*1^+/+^* platelets. Alpha1 deficiency also did not affect collagen-related peptide (CRP), a specific platelet GPVI agonist, induced platelet activation (Fig. 3B). However, ADP-induced aggregation was significantly reduced in the α*1^+/-^* platelet (**Fig. 3C**), suggesting that α1 may have a crosstalk with ADP receptor, P2Y12 or P2Y1. Alpha1 deficiency did not affect thrombin-induced washed platelet activation (data not shown).

**Fig. 3.**
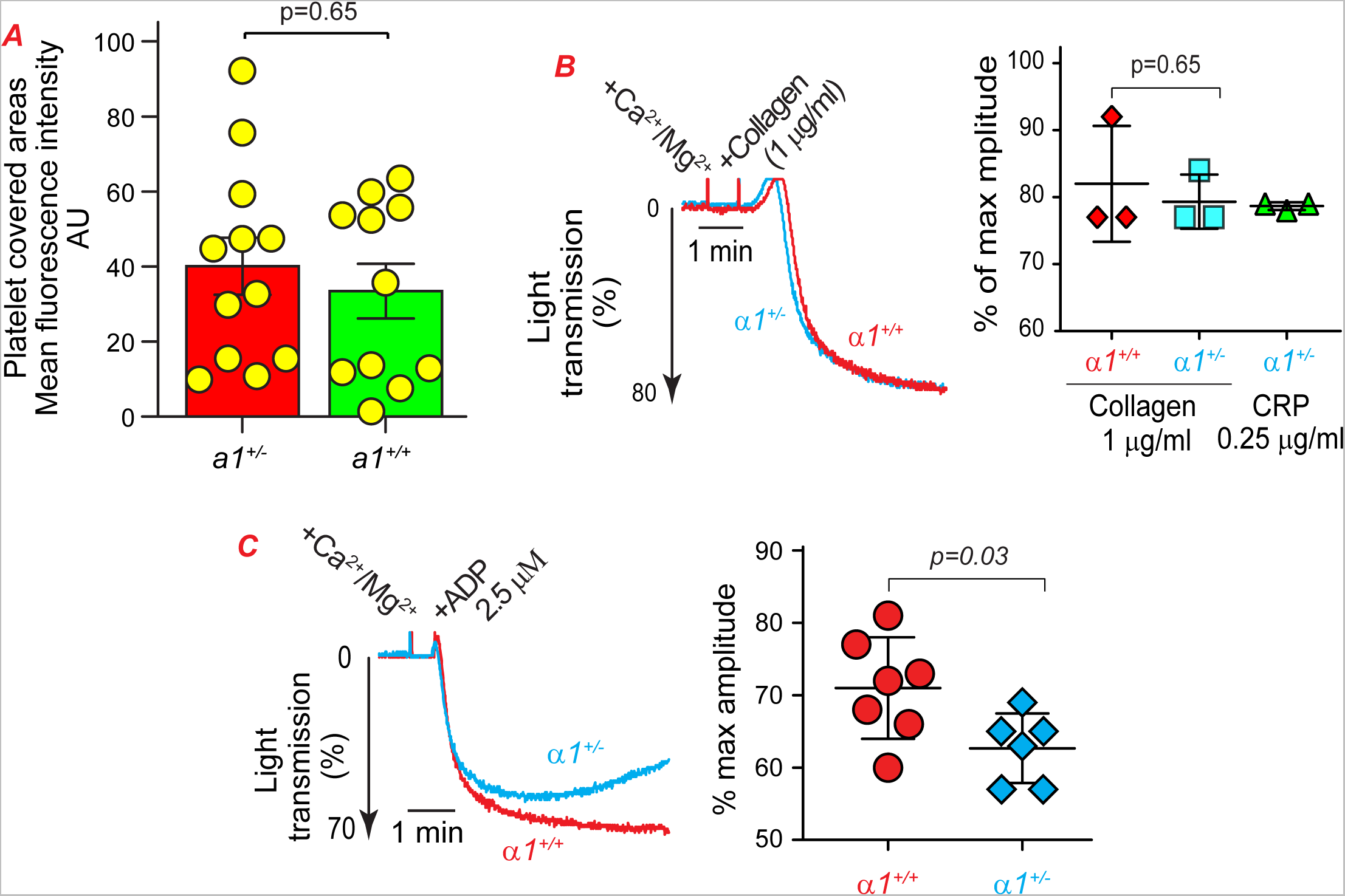
Alpha1 haplodeficiency inhibits ADP-induced platelet aggregation in vitro. ***A.*** Whole blood drawn from *a1^+/+^* and *a1^+/-^* mice was labeled with Rhodamin 6G (50 µg/mL), repleted with CaCl_2_/MgCl_2_ at a final concentration of 1 mM, and then perfused through the collagen (100 µg/mL)-coated Cellix flow chamber at a shear of 60 Dyn/cm² for 3 minutes. The chambers were washed with PBS and then images were taken at predesignated marker positions. Fluorescence areas, representing areas covered by platelets, were quantified by ImageJ and were analyzed with a Student’s t-test. ***B & C.*** Whole blood was drawn from *a1^+/+^*and *a1^+/-^* mice using 0.109 M sodium citrate as an anticoagulant, and platelet-rich plasma (PRP) was prepared by centrifugation. The PRP was repleted with CaCl_2_/MgCl_2_ at a final concentration of 1 mM, and then collagen- and CRP-induced (B) as well as ADP (2.5 µM)-induced platelet aggregation (C) assay was conducted. Student’s *t*-test was used.

### Alpha1 haplodeficiency reduces ADP-stimulated AKT activation and α1 complexes with P2Y12

Both P2Y12 and α1 mediated signaling activates PI3K/AKT^42,55^. We thus treated α*1^+/-^* and α*1^+/+^* platelets with ADP for different durations and examined the phosphorylation of AKT as a marker of P2Y12 signaling activation. As shown in **Fig. 4A**, although ADP activates AKT in both cells, the rate of AKT phosphorylation was dramatically reduced in α*1^+/-^* platelets. These data further demonstrate that NKA α1 affects platelet function, which is probably mediated by the regulation of P2Y12 signaling activation.

**Fig. 4.**
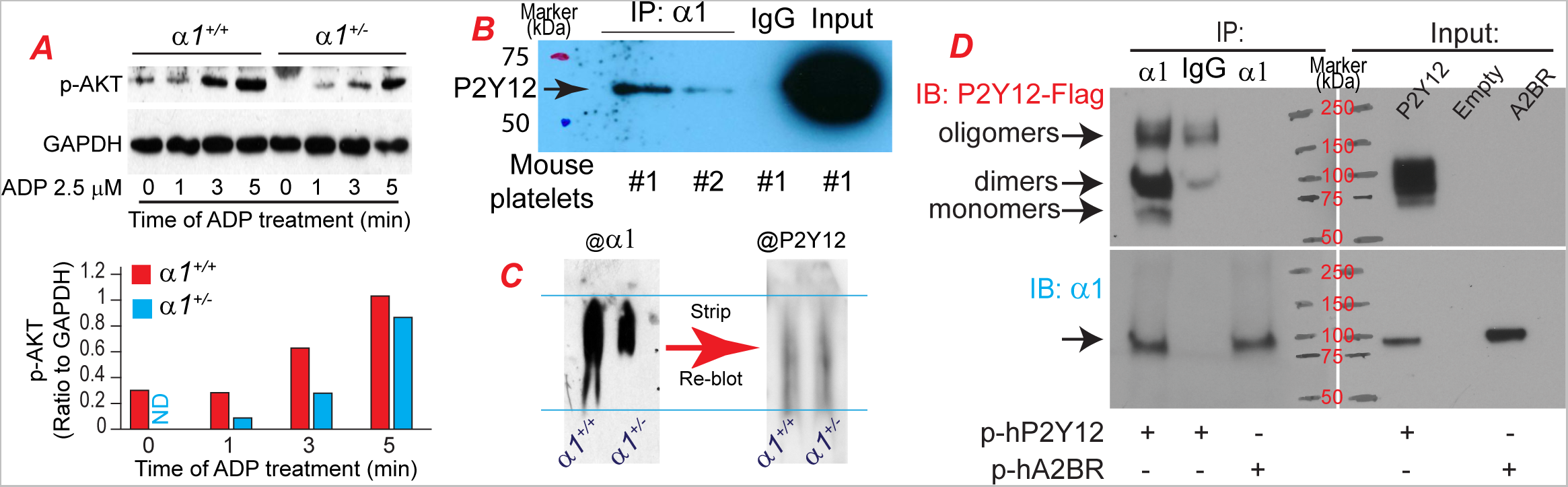
Alpha 1 haplodeficiency inhibits ADP-induced AKT activation in platelets, which may be mediated by the direct binding to P2Y12. *A.* α*1^+/-^* and α*1^+/+^* platelets in PRP were pooled from five mice from the same strain, and platelet concentration was adjusted to 2.5E+08/mL with PPP. The PRP was divided into 4 equal parts, repleted with MgCl_2_/CaCl_2_ to a final concentration of 1 mM, and then treated with 2.5 µM ADP for the indicated periods with rotation. The reaction was stopped by adding an EDTA/PGE1 cocktail containing 500 ng/mL PGE1 and 2 mM EDTA. Platelets were pelleted by a quick spin at 13,000 rpm for 15 seconds, lysed in RIPA buffer, and used for western blot assay of AKT activation. GAPDH was blotted as a loading control. The blot represents two repeats. *B.* WT platelets pooled from 6 mice were lysed and used for IP of NKA α1 (with ab2872 antibody, 2 µg antibody/1 mg total protein), and then IB for P2Y12 was conducted. IP was also conducted with #1 WT sample pool with normal mouse IgG (2 µg normal IgG/1 mg total protein) as control. *C.* Platelet lysates prepared from α1^+/-^ and α*1^+/+^* mice were subjected to the Blue-Native PAGE assay. The membrane was blotted for α1 first, stripped, and re-blotted for P2Y12. The image represents two repeats. *D.* COS-7 cells were transiently transfected with P2Y12-tango plasmid (p-hP2Y12, addgene, #66471), or pEYFP-N1-A2BR (p-hA2BR, addgene, #37202) for 36 h and then cell lysates were used for IP α1 (with ab2872) and immunoblotted for P2Y12. The membrane was stripped and re-blotted for α1 and A2BR (no band was detected in the IP and data was not shown).

To understand how α1 cross talks with ADP receptors, we examined whether α1 complexes with P2Y12 in mouse platelets. As shown in **Fig. 4B**, we found that IP of α1 pulled down P2Y12, but not P2Y1 (data not shown). These findings suggest that α1 may influence platelet function through its direct binding with P2Y12. To further validate this finding, we conducted a Blue-Native PAGE assay using platelets isolated from the α*1^+/-^* or α*1^+/+^* mice. These data also indicated that α1 forms a complex with P2Y12 (**Fig. 4C**). To determine if α1 binding to P2Y12 is a universal feature of these two proteins, we transfected plasmid vector encoding human P2Y12 or adenosine A2B receptor (A2BR) into COS-7 cells, a green monkey kidney fibroblast-like cell line, and used the cell lysates to repeat the IP-IB assay. The green monkey’s α1 (XP_007975629.1) is completely conserved to the human α1 (NP_000692.2) (Data not shown). Both P2Y12 and A2BR are purinergic GPCRs^56^. Under this condition (37 °C, 30 minutes incubation of samples with sample buffer), we found that P2Y12 presents in multiple forms, including monomer, dimer, and multimer. Alpha 1 binds to the P2Y12 dimer the most, and to the monomer the least (**Fig. 4D**). Alpha 1 does not bind to A2BR (data not shown). Taken together, these data suggest that the binding of α1 to P2Y12 could be either P2Y12 specific or depending on the specific structure of these two proteins.

### NKA α1 binds to leucine-glycine-leucine (LGL)-containing platelet GPCRs

The binding of PAR4 and P2Y12 in human platelets is mediated by the peptide leucine-glycine-leucine (LGL)^48^. This LGL motif is conserved in mouse P2Y12 (amino acid 121-123). Using a site-directed mutagenic approach, we designed a mutant variant of mouse P2Y12 and switched the LGL motif to serine-phenylalanine-threonine (SFT)^48^. As shown in **Fig. 5A**, the mutation of LGL to SFT in mouse P2Y12 significantly reduced its affinity to α1.

**Fig. 5.**
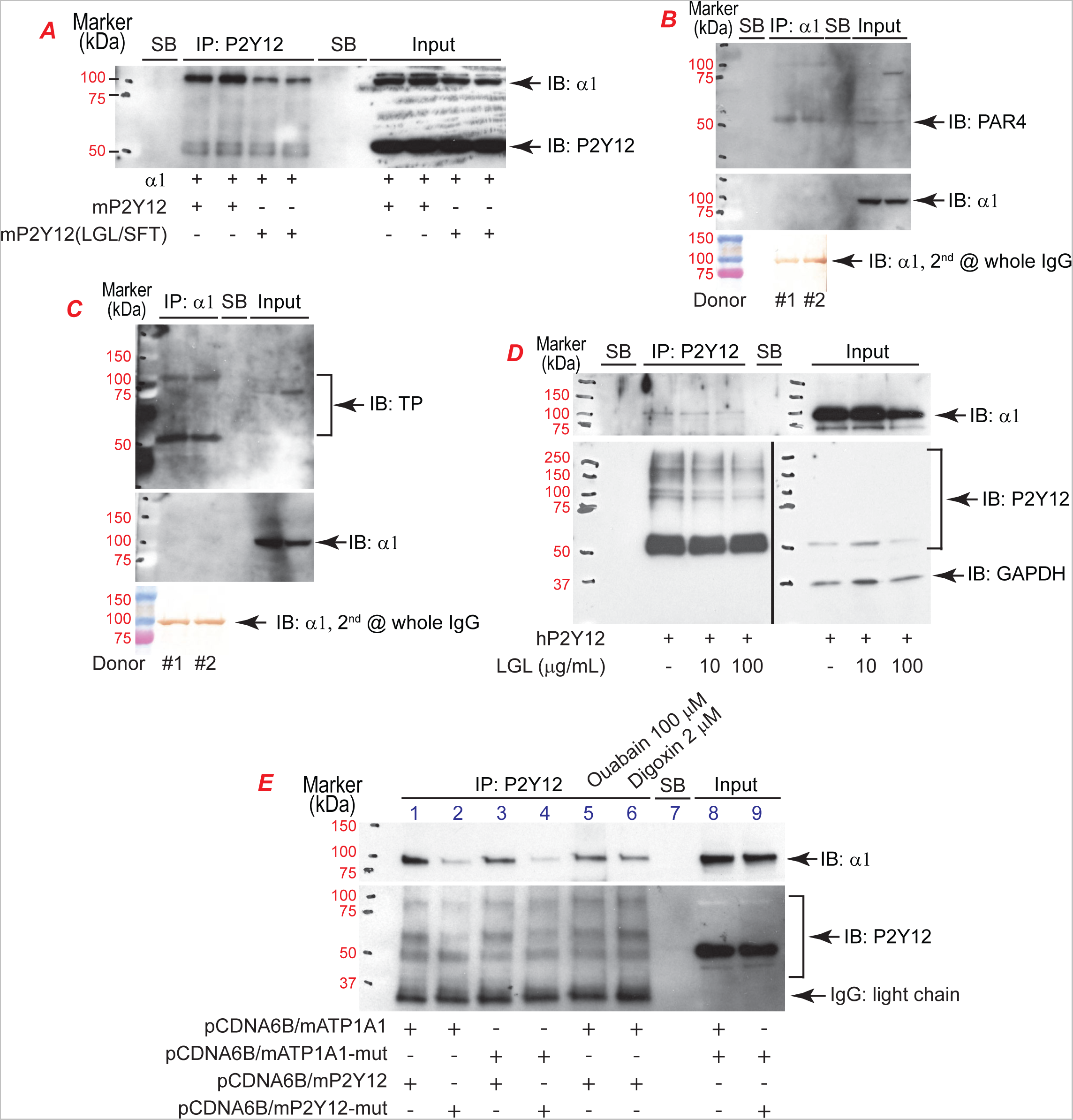
Leucine-Glycine-Leucine (LGL) motif mediates the binding of NKA α1 and certain platelet GPCRs. ***A.*** COS-7 cells were transiently transfected with plasmids encoding mouse NKA α1 and WT or LGL>SFT mutated mouse P2Y12 for 36 hours and then cells were lysed and used for the IP-IB assay. ***B & C.*** Human platelet lysates prepared from two individuals were used for IP of α1 (with ab2872) and then IB for PAR4 and TP. The membranes were stripped and reblotted for α1 (ab76020). HRP-conjugated anti-rabbit IgG light chain antibody was used for the detection of PAR4, TP, and α1 (middle blot) signals. Due to the weak signal of α1 in the IP, the membranes were re-blotted using the HRP-conjugated anti-rabbit whole IgG, and then the α1 signal was visualized by DAB staining. ***D.*** COS-7 cells were transiently transfected with plasmids encoding human P2Y12 for 24 hours. The cells were then treated with LGL at the indicated concentrations for an additional 16 hours. Cells were lysed and used for the co-IP assay. GAPDH was blotted for loading control of input. ***E.*** COS-7 cells were transiently transfected with plasmids encoding mouse α1 and P2Y12, either WT or LGL(LGI)>SFT mutated genes, in different combinations and then cells were lysed and used for the IP-IB assay. Two dishes of cells transfected with mouse WT α1 and P2Y12 were treated with 100 µM Ouabain or 2 µM Digoxin, respectively, for 1 hour before being harvested for IP-IB assay. SB indicates sample buffer only was loaded in the corresponding lane.

Consequently, we analyzed the amino acid sequence of certain conventional human platelet GPCRs, including PAR1, PAR4, P2Y1, P2Y12, TP, PTGER3, PI2R (IP), ADRA2A, and 5HT-2A. We found that in addition to P2Y12 (115-117), PAR4 (194-196), PTGER3 (237-239), TP (76-78 and 163-165), and PI2R (IP) (82-84 and 156-158) also contain LGL motif(s). We thus further examined whether α1 also binds to PAR4 and TP, both mediate platelet activation. To this end, human platelet lysates were used for the co-IP assays. As shown in **Fig. 5B** and **5C**, pulling down α1 also pulled down PAR4 and TP. Again, P2Y1 was not pulled down by α1 (Data not shown). These data further demonstrate that α1 binds to LGL-containing platelet GPCRs, independent of species, and thus may affect these GPCRs-mediated platelet activation.

We next examined if LGL peptide could serve as a decoy and block the complex formation between α1 and LGL-containing platelet GPCRs. COS-7 cells were transfected with human P2Y12 for 24 hours, then treated with LGL in the complete cell culture media in a final concentration of 0, 10, and 100 µg/mL, and cultured for 16 hours. The cells were harvested and lysates were used for co-IP assay. As shown in **Fig. 5D**, LGL treatment significantly attenuated the binding of α1 to P2Y12. LGL treatment did not cause cell toxicity (**Supplemental Figure 2A**) and also did not affect the expression of α1 (**Supplemental Figure 2B**).

Since human α1 also contains an LGL motif, which is conserved in mouse α1 as LGI, we mutated mouse LGI to SFT, and then examined its affinity to mouse P2Y12. As shown in **Fig. 5E**, LGI>SFT mutation in mouse α1 also attenuated the association of P2Y12 and α1 (lane 3 vs lane 1). Co-transfection of the LGI(LGL)>SFT mutant α1 and P2Y12 further diminished their binding (lane 4 vs lane 1 to 3). Treating the cells co-transfected with mouse α1 and P2Y12 with ouabain (lane 5) or digoxin (lane 6) also reduced the association of α1 and P2Y12.

### A therapeutic dose of digoxin enhanced α1 and P2Y12 dimer and oligomer formation

To clarify how CTS affects platelet function and thrombosis, we collected blood from HF patients on digoxin and compared their platelet activity with healthy donors. As shown in **Fig. 6A**, we found that digoxin treatment did not enhance 10 µM ADP-stimulated platelet aggregation. Rather, it tended to reduce platelet aggregation.

**Fig. 6.**
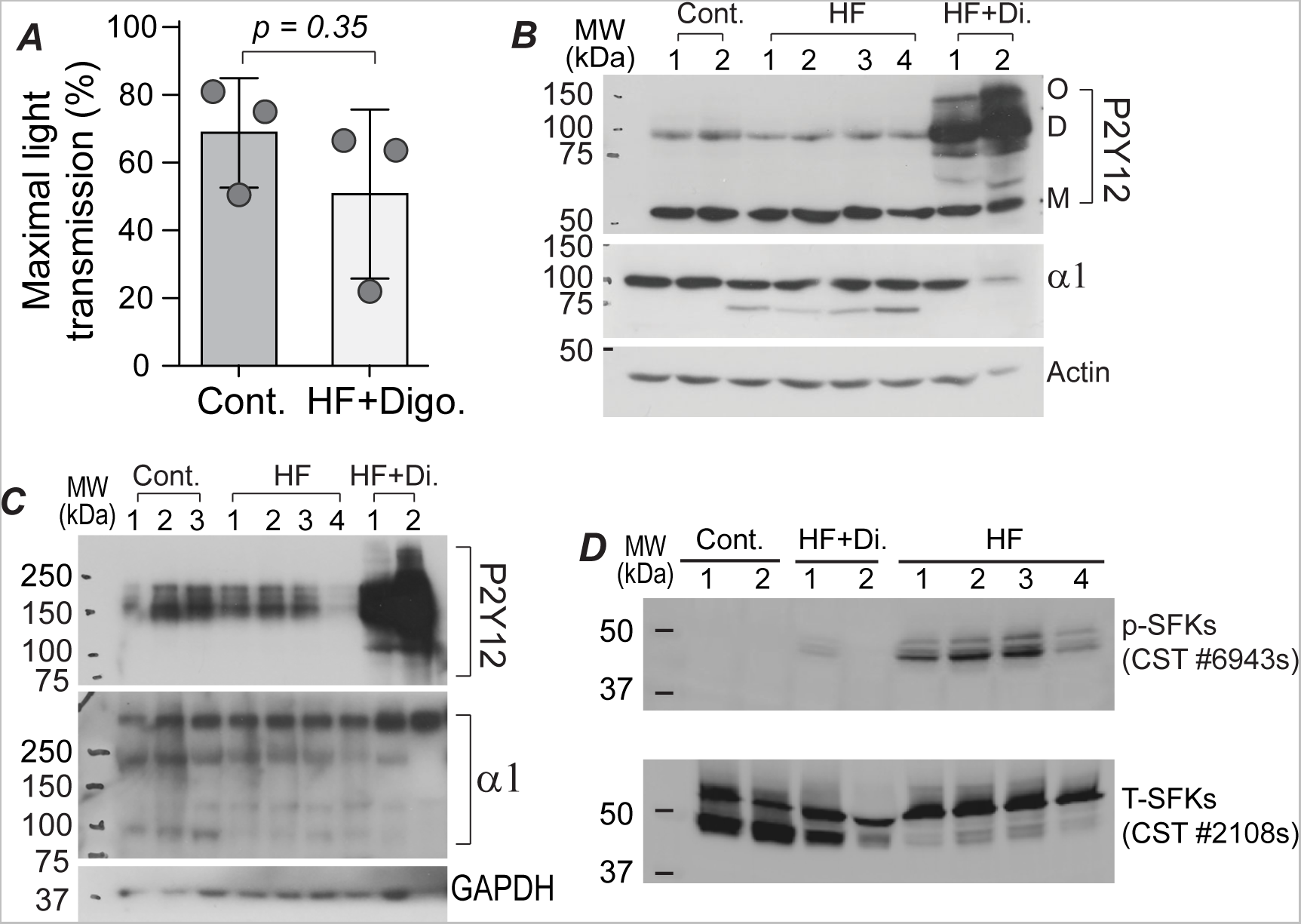
Therapeutic dose of digoxin (Di.) enhanced α1 and P2Y12 dimerization but inhibited the activity of Src Family kinases (SFKs). Whole blood was drawn from healthy donors (Cont.), heart failure (HF) patients, or HF patients receiving therapeutic doses of digoxin (HF+Di.) and platelets were isolated by centrifugation. ***A.*** Platelet-rich-plasma was prepared and used for aggregation assay induced by 10 µM ADP. ***B & C.*** Platelets were washed two times with sodium citrate/saline solution (1:9 ratio) in the presence of 100 ng/mL PGE1 and then lysed in RIPA buffer. The sample concentration was adjusted to 2 µg/µL with RIPA buffer. Samples were mixed with an equal volume of 2X Laemmli sample buffer in the presence ***(B)*** or absence ***(C)*** of 2-mercaptoethanol and incubated at 37 °C for 30 min. Forty micrograms of total proteins were used for the western blot assay of the proteins indicated. O: Oligomer; D: Dimer; M: monomer. ***D.*** Samples used in B were used for western blot assay of SFK activity, detected by phosphor-Src antibody. The membrane was stripped and reblotted with anti-total Src antibody as loading control.

We thus examined α1 and P2Y12 expressions in these platelets and used platelets harvested from HF patients without digoxin treatment as controls. As shown in **Fig. 6B**, under a reduced condition, we found that both α1 and dimeric P2Y12 were decreased in HF platelets. Interestingly, P2Y12 dimer and oligomer were dramatically increased in platelets of HF patients receiving digoxin under both reduced and non-reduced conditions (**Fig. 6B** & **6C**). One band above 250 kDa was found in the non-reduced condition for α1 blot, which could be either tetrameric α1 or α1β3 diprotomer^57^ or other α1-associated protein-protein complex, was increased in HF platelets, and markedly increased in platelets of HF patients on digoxin.

To further understand the platelet activity under these conditions, we measured the activity of Src-family kinases (SFKs), as SFKs are known to play important roles in platelet activation^58^. As shown in **Fig. 6D**, the activated (phosphorylated) SFKs were significantly increased in platelets harvested from the HF patients, but not in control platelets. Digoxin treatment dramatically reduced the active SFKs.

### Alpha1 haplodeficiency significantly lowered the effective dose of clopidogrel in inhibiting thrombosis and low-dose CTS inhibited thrombosis in mice and dose-dependently attenuated ADP-induced human platelet aggregation

We previously demonstrated that treating WT mice with 1 mg/kg of clopidogrel has uncertain effects on inhibiting thrombosis in vivo^42^. We thus treated α*1*^+/+^ and α*1*^+/-^ mice with 0.5 or 1 mg/kg of clopidogrel by gavage feeding for one week and subsequently evaluated thrombosis in these mice using the 10% FeCl_3_-induced carotid artery injury thrombosis model, as under this condition, α1 haplodeficiency did not achieve an antithrombotic effect. Treating α*1*^+/-^ mice with 0.5 mg/kg clopidogrel did not significantly prolong the time to form occlusive thrombi (**Fig. 7A**). However, treating mice with 1 mg/kg clopidogrel for 1 week significantly prolonged the time to thrombosis in the α*1*^+/-^ mice.

**Fig. 7.**
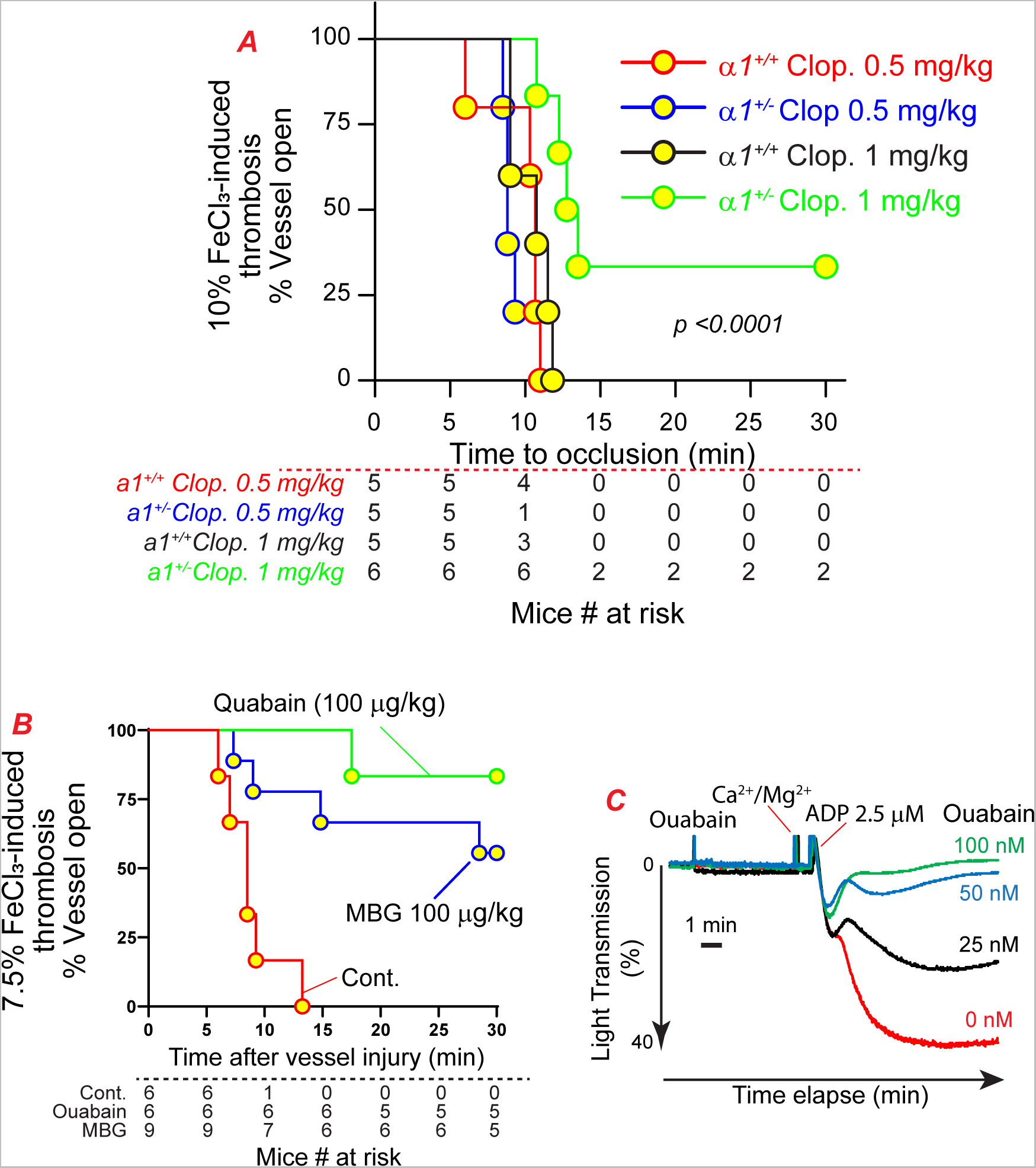
Alpha1 haplodeficiency lowers the effective dose of clopidogrel needed to inhibit thrombosis and low-dose CTS inhibits thrombosis in mice and dose-dependently attenuates ADP-induced human platelet aggregation. *A.* α*1^+/-^*and α*1^+/+^* mice were gavage-fed clopidogrel at a dose of 0.5 and 1 mg/kg for one week and then subjected to the 10% FeCl_3_-injury induced thrombosis model. *B.* WT mice were treated with ouabain (100 µg/kg) or MBG (100 µg/kg), by bolus intraperitoneal injection, and then subjected to the in vivo thrombosis study 18 h later. *C.* Human PRP was pre-treated with ouabain in the indicated concentrations for 5 min, and then aggregation was initiated with 2.5 µM ADP. The logrank test was used for the statistical analyses in panels A and B.

To determine if CTS treatment mimics α*1^+/-^* mice, we treated WT mice with ouabain (100 µg/Kg body weight) via intraperitoneal injection, and a thrombosis study was conducted 18 h later. As shown in **Fig. 7B**, one dose of ouabain treatment significantly inhibited thrombosis in mice (*p = 0.005* vs cont.), which is in line with our findings in α*1^+/-^* mice.

The CTS MBG, a member of the bufadienolide family, also inhibits NKA function. MBG shares a conserved structural core with the cardenolides ouabain and digoxin^59^, and the binding site of ouabain interacts with MBG^60^. Intraperitoneal injection of 560 µg/kg of MBG significantly attenuated zymosan-induced inflammation^50^. We treated WT mice with a low dose of MBG (100 µg/kg) overnight and then examined its impact on thrombosis. As shown with the blue line in **Fig. 7B**, low-dose MBG also significantly inhibited thrombosis in mice (*p = 0.003* vs cont.).

Since rodent α1 has ∼100-1000-fold less responsiveness to ouabain than the α1 in humans and other mammals^18^, we examined whether the different observations between our study and the published data are due to the difference of species. We pre-treated human platelet-rich plasma prepared from healthy donors with different concentrations of ouabain for 5 min and then initiated the aggregation with 2.5 µM ADP. As shown in **Fig. 7C**, in certain platelet donors, ouabain dose-dependently inhibited ADP-induced platelet aggregation. Of note, only two of the six healthy donors (33.3%) examined showed this type of response. We did not find ouabain to potentiate or inhibit ADP-induced platelet activation in the other four cases tested (66.7%), suggesting that different individuals may have different sensitivity to ouabain. Taken together, these data further suggest that α1 plays a role in platelet activation, and a link between α1 and the observation that low-dose CTS inhibits platelet activation and thrombosis.

## Discussion

This study represents, to the best of our knowledge, the first systemic study on the role of NKA in thrombosis using genetically modified mice and several CTS using both in vivo and in vitro approaches. We have uncovered novel mechanisms that mediate the regulation of platelet activation and thrombosis. The major findings include 1) α1 haplodeficiency results in a significant anti-thrombotic effect in male mice without disturbing hemostasis; 2) α1 forms complexes with P2Y12, PAR4, and TP, all of which are GPCRs containing at least one LGL motif. Haplodeficiency of α1 reduces ADP-stimulated AKT activation in platelets and attenuates ADP-induced platelet aggregation; 3) haplodeficiency of α1 significantly enhances the efficacy of clopidogrel in inhibiting thrombosis in vivo; 4) Low-dose CTS inhibit thrombosis in mice and attenuate ADP-induced platelet aggregation in certain human platelets.

A proteomics study has revealed that human platelets harbor approximately 2,000 copies of NKA α1, 500 copies of α4, and 2,500 copies of β3 subunits^20^. Notably, the α4 subunit is exclusive to sperm, sharing 78% identity with α1^61^, and it was not identified in a previous human platelet study^21^ and a mouse platelet proteomics study^22^. Consequently, the platelet NKA is presumed to be comprised of α1 and β3 subunits. The ouabain-sensitive α2 and α3 isoforms were also not identified in platelets^20–22^. Of note, the Cre-Loxp strategy was unable to produce PF4-Cre^+^/α*1^f/f^*mice (data not shown), suggesting that complete α1 deficiency in platelets results in embryonic lethality in mice. These findings support our hypothesis that α1 is the sole NKA α isoform in platelets.

As mentioned in the Introduction, the role of CTS on human platelet activation is under debate^26^. Scattered studies suggested that inhibiting NKA pump function with ouabain enhances human platelet aggregation in vitro^28–30^, mediated by the increase in intracellular Ca^2+^-evoked exposure of phosphatidylserine^47^. Notably, these studies employed very high concentrations of ouabain (10-200 μM)^28,29^. Given that the IC50 of ouabain is less than three-digit nanomolar for two human cancer cell lines^62^, the impact of ouabain on platelet function at high doses needs to be further studied and compared with the effect of ouabain at low doses. We found that treating WT mice with 100 ug/kg ouabain via intraperitoneal injection overnight significantly inhibited thrombosis in vivo. This observation highlights the need for a comprehensive understanding of the dose-dependent effects of ouabain and other CTS on platelet function and thrombosis.

Accumulated studies suggest that GPCRs can form homo- or hetero-dimers and even higher-order oligomers, with dimerization/oligomerization being necessary for signaling initiation^63,64^. As mentioned above, the binding between PAR4 and P2Y12 in human platelets is mediated by LGL^48^. In this study, we have made a novel discovery: the LGL motif also facilitates the binding between NKA α1 and several platelet GPCRs, including P2Y12, PAR4, and TP. These bindings may be through a two-domain interaction, as the LGL motif is present in both binding partners. These findings were solidified by introducing mutations LGL(I)>SFT in both α1 and P2Y12, as well as using a custom-generated LGL peptide as a decoy. Interestingly, the LGL motif has been reported as a foodborne peptide found in Parma Dry-Cured Ham. This peptide has been shown to have a strong inhibitory effect on angiotensin I converting enzyme (ACE)^65^, which contains three LGL motifs (based on NP_000780.1). Additionally, Spanish dry-cured ham, processed similarly to the parma dry-cured ham, has been reported to attenuate platelet and monocyte activation^66^, although LGL was not mentioned in that study. Furthermore, treating COS-7 cells with LGL peptide did not show any cell toxicity effect, even when a concentration of 100 µg/mL dose was used. Taken together, these findings suggest that targeting α1 and platelet GPCRs with the LGL peptide could represent a novel mechanism-based antithrombotic strategy.

Ouabain and digoxin have been clinically used in the setting of HF as well as atrial fibrillation. Clinically use of digoxin has been associated with increased platelet and endothelial cell activation in patients with nonvalvular atrial fibrillation^31^, a condition often seen in patients with HF. In our study, we found that therapeutic doses of digoxin enhanced the formation of dimer and oligomer involving α1 and P2Y12 (maybe PAR4 and TP too). In HF platelets, a 75-80 kDa band was detected by the anti-α1 antibody. However, it is absent in platelets from normal blood donors and HF patients receiving digoxin. A recent study indicated that the binding of ouabain to α1 enhances α1 trypsinolysis^67^. The increase in endogenous digitalis-like compounds has been reported in HF patients, with circulating levels ranging from 1-9 nM^68,69^. However, some groups have challenged this concept, stating that “No endogenous ouabain is detectable in human plasma”^70^. Nevertheless, the presence of endogenous ouabain or other factors could partially explain our findings, as low-dose ouabain can activate c-Src^49^, and digoxin-ouabain antagonism can diminish c-Src activation^71^. Whether the 75-80 kDa peptide found in HF platelets is due to trypsinolysis of α1 and whether it affects platelet function will be an interesting aspect to pursue in future studies. Taken together, these findings shed light on exploring a novel mechanism regarding digoxin-associated thrombosis in HF. Additional studies using precisely manipulated doses of digoxin and other CTS and testing their effect on platelet activation as well as in thrombosis are needed to clarify the clinical significance of these dimerizations.

Current antiplatelet and antithrombotic medications typically require chronic dosing to achieve effectiveness, and they are associated with bleeding side effects. Since α*1* deficiency does not affect tail-bleeding time, and a single dose of ouabain or MBG administration significantly inhibits thrombosis in vivo, this study suggests that α1 could be a target for dual antiplatelet therapy. This is supported by the observation that an uncertain dose of clopidogrel treatment significantly inhibited thrombosis in the α*1^+/-^* mice but not in the α*1^+/+^* mice. Studies have shown that a cell only needs 20-30% of its maximal NKA pump expression to maintain normal intracellular Na^+^/K^+^ gradients^72,73^. Therefore, inhibition of platelet NKA α1 can be a safe mechanism of antithrombotic therapy, either administered alone or in combination with other medications.

In summary, this study is the first to systemically investigate the role of NKA α1 in platelet activation and thrombosis. We found that α1 binds to LGL motif-containing platelet GPCRs, including P2Y12, PAR, TP, and potentially others. NKA α1 plays an essential role in fine-turning platelet activation. Targeting NKA α1 could be a novel strategy for antiplatelet and antithrombotic therapy.

## Acknowledgments

The authors thank Mrs. Melissa Marcum for consending heart failure patients and helping prepare the IRB protocols. The authors thank Dr. Jerry Lingrel for providing the α*1^+/-^* mice.

## Authorship Contributions

Study design and drafting manuscript:: OL, HY, SP, and WL.

Data Collection: RR, YH, AD, RR, RA, OO, FB, JL, DK, and WL.

Critical Revisions: OF, DK, ET.

Final approval for publication: SP and WL.

## Abbreviations

NKA: Sodium-Potassium ATPase
HF: Heart failure
ADP: Adenosine diphosphate
TXA2: Thromboxane A2
GPCR: G protein-coupled receptors
TP: TXA2 receptor
CTS: Cardiotonic steroids
WT: Wild-type
CRP: Collagen-related peptide
PVDF: Polyvinylidene difluoride
DTT: Dithiothreitol
FeCl_3_: Ferric chloride
PRP: Platelet-rich plasma
PPP: Platelet-poor plasma
CPM: Counts per minute
RIPA: Radioimmunoprecipitation assay
BM: Bone marrow
DMEM: Dulbecco’s Modified Eagle Medium
FCS: Fetal Calf Serum
IP: Immunoprecipitation
IB: Immunoblotting
DDM: Dodecyl-β-d-maltoside
MBG: Marinobufagenin
DAB: 3, 3’-diaminobenzidine
LGL: Leucine-glycine-leucine
SFK: Src-family kinase

