## Supplementary figures and images for "Sodium/Potassium ATPase Alpha 1 Subunit Fine-tunes Platelet GPCR Signaling Function and is Essential for Thrombosis"

### Supplemental Figure 1

Supplemental Figure 1

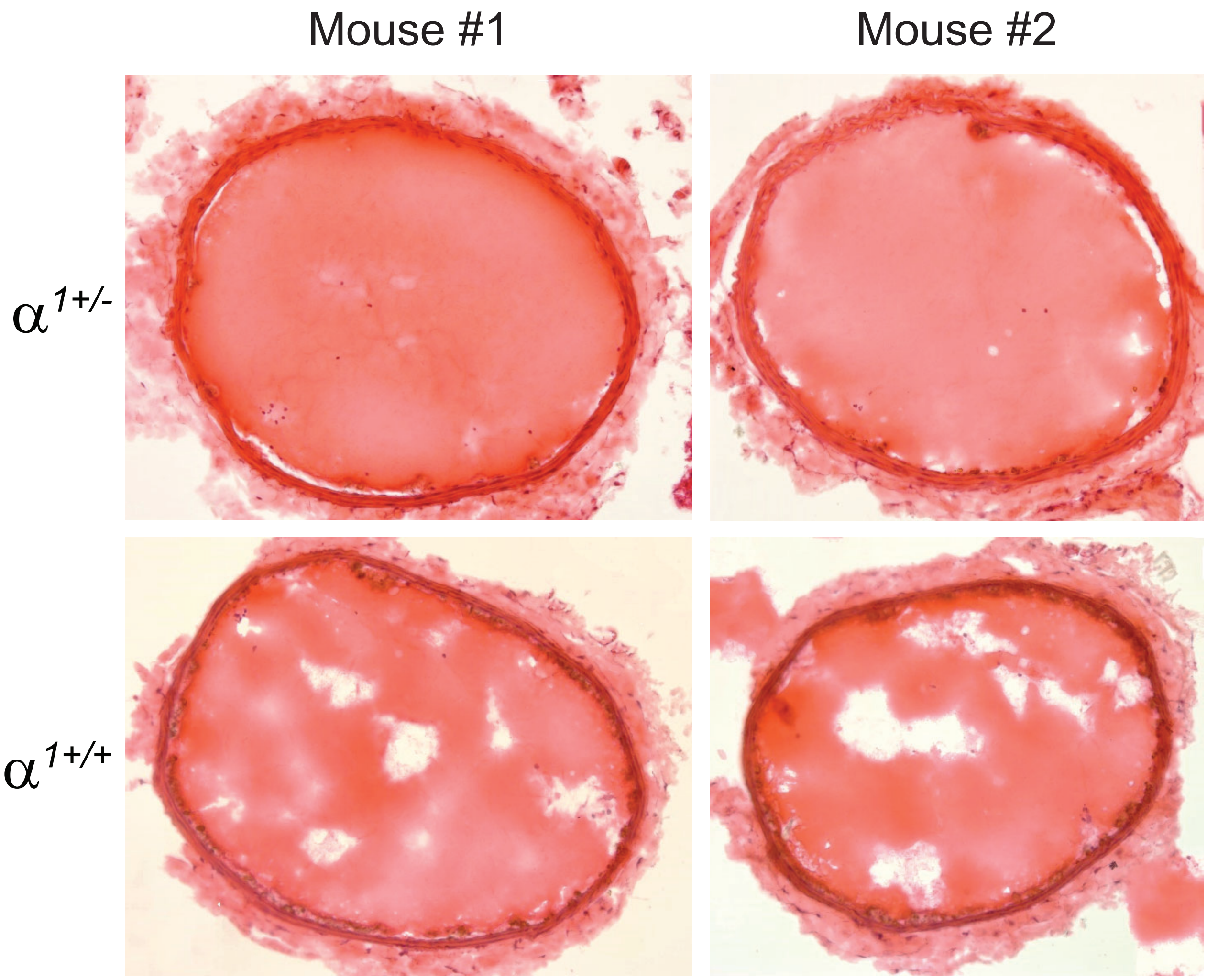

### Supplemental Figure 2

Supplemental Figure 2

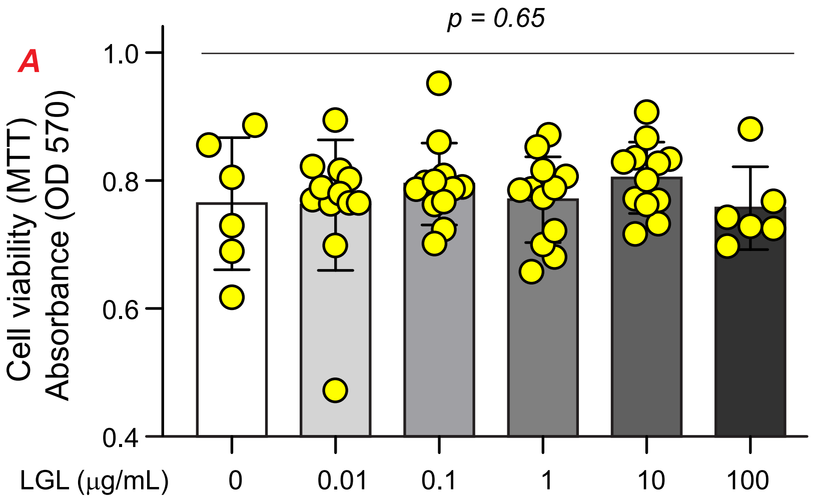


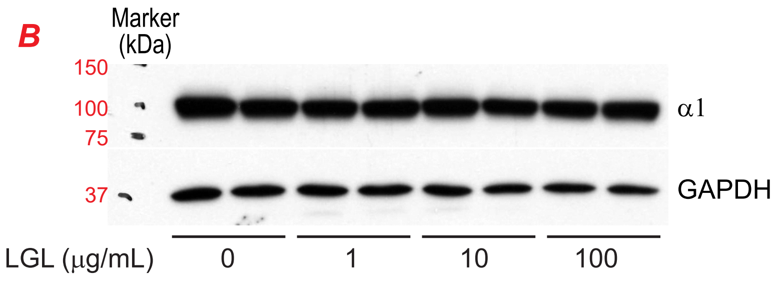
